# Brain, immune system and selenium: a starting point for a new diagnostic marker for Alzheimer’s disease?

**DOI:** 10.1101/239293

**Authors:** Cesare Achilli, Annarita Ciana, Giampaolo Minetti

**Author notes:** Corresponding author: Giampaolo Minetti.

## Abstract

The clinical diagnosis of Alzheimer’s disease (AD) is based primarily on neuropsychological tests, which assess the involutive damage, and imaging techniques that evaluate morphologic changes in the brain. The currently available diagnostic tests do not show complete specificity and do not permit an accurate differentiation between AD and other forms of senile dementia. The correlation of these tests with laboratory investigations based on biochemical parameters could increase the certainty of the diagnosis. In recent years, several biochemical markers for the diagnosis of AD have been proposed, but in most cases they show a limited specificity and their application is invasive, because it generally requires the sampling of cerebrospinal fluid. Therefore, the use of a peripheral biochemical marker could represent a valuable complement for the diagnosis of this disease.

Several studies have shown a relationship between the neurodegenerative disorders typical of the ageing process, the weakening of the immune system, alterations in the levels of selenium and of the antioxidant selenoenzymes in brain tissues and blood cells, particularly in neutrophil granulocytes. The levels of peripheral selenoenzymes may reveal a promising clinical parameter for helping in the assessment of the pathological condition in AD.

**Highlights:** MsrBl is one of the 25 selenoenzymes expressed in the humans

MsrBl is highly expressed in human circulating neutrophils

The diagnostic markers for Alzheimer’s disease are still insufficiently validated

The impairment of some selenoenzymes is associated with Alzheimer’s disease

Neutrophil MsrBl can be a peripheral marker for the diagnosis of Alzheimer’s disease

## Background

Selenium is an essential element involved in several cellular processes such as immune response, signal transduction, metabolism of thyroid hormones and defence against oxidative damage [1]. Selenium is contained in proteins as selenocysteine, which can be considered as the 21^st^ proteinogenic amino acid. When selenocysteine is present in the active site of the protein, it confers to the enzyme a higher catalytic efficiency compared to cysteine. The most recent studies have revealed that 25 selenoproteins are expressed in humans [2, 3]. Many of them, such as glutathione peroxidase (GPx) and methionine sulfoxide reductase B1 (MsrB1), are oxidoreductase enzymes that contain selenocysteine in the catalytic site and are involved in the regulation of cellular redox processes [4].

Members of the GPx family perform the detoxification of reactive oxygen species (ROS) produced during the aerobic metabolism, through the reduction of organic peroxides using glutathione as the electron donor, and are characterised by different substrate specificity, cellular localization and tissue distribution [5].

During ageing, and in association with neurodegenerative diseases such as Alzheimer’s disease (AD), a significant decrease of GPx activity in whole blood [6], erythrocytes [7–12], neutrophils [7, 13] and plasma [7, 14, 15] was observed. Quite remarkably neutrophils from patients affected by neurodegenerative diseases displayed less than 30% of the GPx activity of neutrophils from healthy individuals [7]. This phenomenon was correlated with the decrease in selenium levels that was observed to occur in plasma [7, 16, 17, 9, 18] and in erythrocytes [7, 18] during ageing and in individuals affected by various neurodegenerative disorders. In a sample of geriatric population, a 25% decrease in selenium plasma levels was observed, along with a 20% decrease in erythrocyte GPx activity [19]. The changes in selenium status that occur during ageing, which can be attributed to defects of assimilation and/or metabolism of this element, may lead to an increased susceptibility to various degenerative diseases typical of the elderly, including AD, which is characterised by oxidative damage to various cellular components [20]. The increased oxidative stress associated with selenium deficiency has been correlated with defects in the regulation of GPx expression [21]. Low selenium levels increase significantly the risk of mortality during senility [22]. This element is therefore considered as an important factor for maintaining a good state of health in the elderly [23]. Components of both the innate and acquired immune systems deteriorate gradually during ageing [24, 25], and this functional impairment further aggravates in the course of AD [26, 27]. The cellular components that guarantee the efficiency of the innate immune response are the phagocytes: macrophages and neutrophil granulocytes, whose task is to engulf the pathogen into a phagosome and degrade it by means of proteolytic enzymes and various ROS, like superoxide, hydrogen peroxide and hypochlorite, produced during the so-called oxidative burst. As ROS can diffuse from the phagosome into the cytosol during the burst, and thus compromise cell function, neutrophils are endowed with powerful antioxidant systems, both enzymatic and non-enzymatic, that act to prevent or repair the oxidative damage [28]. Selenium is essential for the full efficiency of various components of the immune system, both acquired and innate [29, 30]. Several studies conducted in mice and humans have shown that selenium deficiency, as indicated by the loss of GPx activity, leads to a reduction in the phagocytic capacity and in the intensity of the oxidative burst in neutrophils, resulting in decreased bactericidal activity [31, 32], whereas selenium supplementation in the diet restores GPx activity and cell function [32].

Studies in vitro with HL-60 cells (a human promyelocytic leukaemia cell line that can be induced to differentiate into a neutrophilic phenotype) grown in a selenium-deficient medium indicate the complete absence of GPx, whose expression is restored to values comparable to those measured in neutrophils when the medium is supplemented with appropriate amounts of selenium [33, 34]. Another study indicate that the supplementation of selenium in the diet of humans with selenium deficiency associated with neurodegenerative diseases increases both selenium levels and GPx activity in plasma and erythrocytes [7]. These data indicate that GPx expression is very sensitive to the levels of selenium in the organism.

The decrease in neutrophil’s bactericidal capacity appears to be linked to ROS accumulation and to correlate with the loss of GPx activity, resulting in a redox imbalance in the cells, which become therefore more sensitive to oxidative damages. Neutrophils from AD patients were found to contain high concentrations of intracellular ROS under restingesting conditions, pointing to a defect in antioxidant systems of these cells [35].

Human neutrophils are the cell type with the highest selenium concentration, although it has been shown that the levels of GPx activity are lower than those measured in other cell types, such as in kidney and liver. Moreover, neutrophils of mice subjected to a diet supplemented with selenite display, at low selenium concentrations, an increase in bactericidal activity but not in GPx activity, while only at higher concentrations of selenium the two activities increase in parallel [32]. These data suggest that the expression of GPx is particularly dependent on the selenium status of the organism, and, most importantly, that this protein is not the only selenoenzyme that ensures proper neutrophil function.

Only one other selenoenzyme with anti-oxidant function has been characterised in detail in human neutrophils [36]. It is MsrB1, belonging to the family of methionine sulfoxide reductases (MsrA and MsrB), and the only form of Msr containing a catalytically active selenocysteine [37]. The Msrs catalyse the stereospecific reduction of the two diastereoisomers of methionine sulfoxide, which are generated by the oxidation of the sulphide group of methionine, as the free amino acid or when it is present in proteins: [MsrAs reduce (Sc)methionine-(S_S_)sulfoxide, while MsrBs reduce (Sc)methionine-(RS)sulfoxide] [38]. Msrs perform three main tasks: (i) the repair of the oxidative damage suffered by methionine residues, which results in the alteration of function of many proteins, such as the beta-amyloid peptide, in which the oxidation of methionine appears critical for aggregation, neurotoxicity, and the generation of ROS induced by the peptide [39]; (ii) the detoxification of ROS through the cyclic oxidation/reduction of methionine, free or inserted in some proteins, thus preventing the oxidation of other particularly sensitive cellular components [40]; (iii) the regulation of cellular metabolism through changes in the redox state of specific methionine residues present in various enzymes, hormones and plasma proteins [41]. The imbalance in the methionine/methionine sulfoxide ratio can alter the metabolic functions, contributing to the development of diseases related to the impairment of the cellular processes affected [42].

The selenoenzyme MsrB1 is the form predominantly expressed in circulating human neutrophils, but is almost entirely absent in other blood cells [36]. MsrB1, like GPx, would be a component of the selenium-dependent antioxidant machinery that has evolved in neutrophils to counteract the ROS produced in high concentrations during the immune response.

## Hypothesis

We propose this hypothesis: the selenoenzyme MsrB1 from neutrophil granulocytes could represent a more specific indicator of the selenium status in the organism and in particular a new peripheral diagnostic marker for AD, alternative to, or in combination with those proposed, which are typically invasive since they involve the analysis of cerebrospinal fluid, and display low specificity [43, 44].

## Results and Discussion

Concerning the hypothesis presented here, MsrB1 activity was analysed in neutrophil samples from healthy donors and patients affected by AD associated with diversified diseases typical of the elderly. A limited cohort of patients was studied, not allowing statistical elaboration, therefore the present study should be considered as a preliminary investigation. Yet, this pilot study showed that the patients could be grouped in two populations: one group with MsrB1 specific activity similar to that of healthy donors, and one group with approximately the 50% of the specific activity of healthy donors (Figure 1).

**Figure 1.**
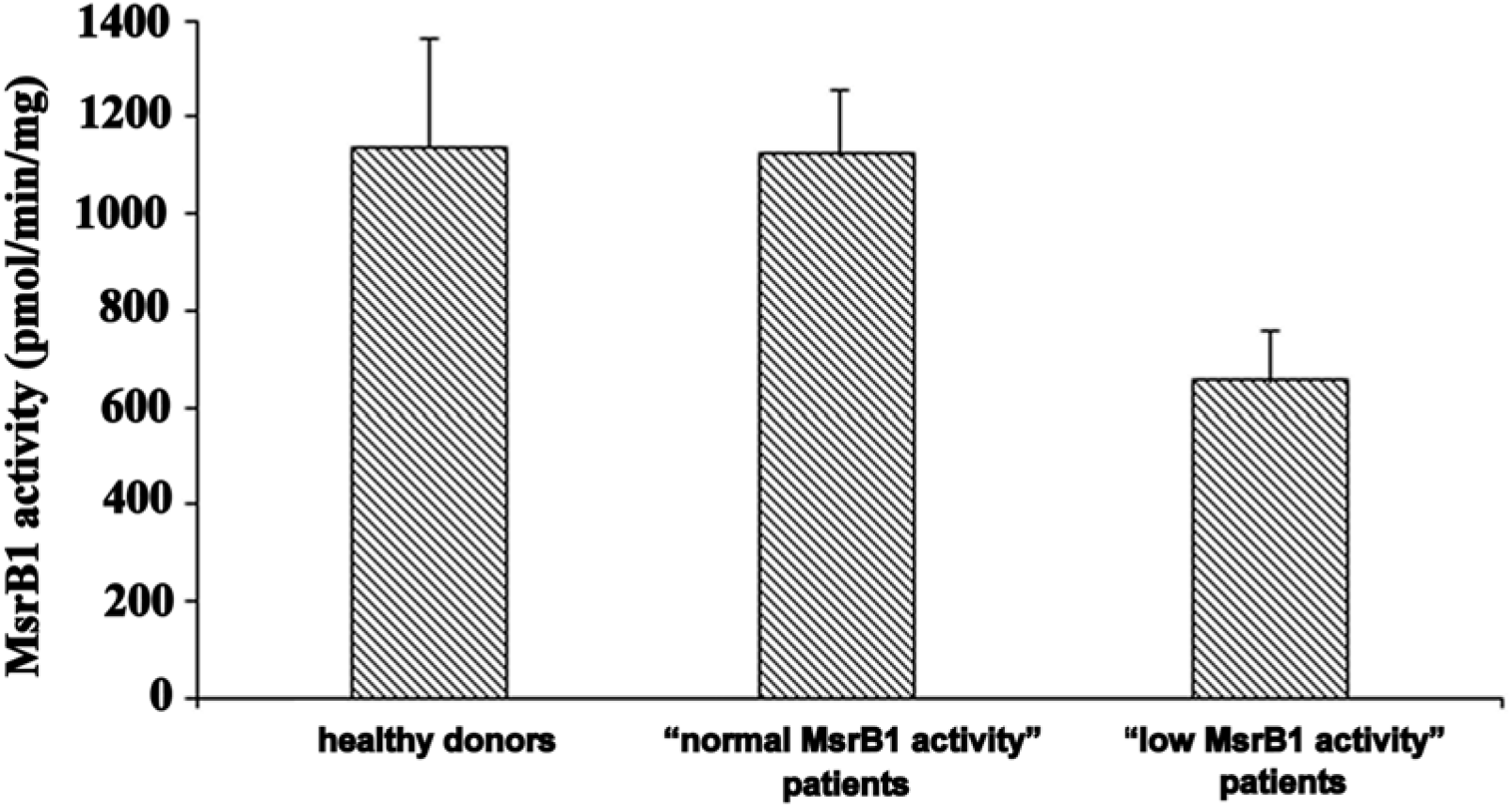
Specific activity of MsrB1 (pmol of dabsyl-methionine produced per min per mg total protein) detected in neutrophils from: healthy donors (n=8), “normal MsrB1 activity” patients (n=8), and “low MsrB1 activity” patients (n=5).

The novel finding of decreased MsrB1 activity with ageing presented here is consistent with literature data about the decrease in antioxidant defence, and particularly of selenoenzymes, associated with age-related neurodegenerative pathologies typical of the elderly. Moreover, particularly remarkable is a post-mortem observation that showed a significant decrease in Msr activity (MsrA plus MsrB) in various areas of the brain in subjects affected by AD [45]. The alterations observed in the immune system of patients suffering from neurodegenerative disorders, particularly AD, suggest a correlation between the two events, supporting the hypothesis of a systemic nature of this disease [26, 46]. The observation of a concomitant impairment of Msr activity in brain tissues and peripheral neutrophils, in association with AD, seem to corroborate this hypothesis.

Moreover, the increase in the intracellular content of oxidized proteins, especially at the level of methionine residues, which occurs during ageing, has been correlated with the onset of degenerative diseases typical of advanced age [42, 47, 48]. The limited capacity of reduction of methionine sulfoxide by Msr enzymes, including the selenoenzyme MsrB1, renders the defence systems against oxidative stress inadequate, thus favouring the onset of degenerative diseases, like the Alzheimer’s, which are characterised by accumulation of oxidized proteins [42, 47].

In light of the preliminary evidence presented here, the idea of a correlation between AD, partial impairment of the immune system and decreased MsrB1 activity should be further investigated. It is also to be clarified whether the decline MsrB1 activity observed in some AD patients was due to a decrease in protein expression or to an impairment of the catalytic activity. Although MsrB1 was discovered in 1999 [38], it is still poorly studied and it has not yet been considered as a possible indicator of the selenium status of the organism [49]. Therefore, we deem it of particular interest to explore the potential use of MsrB1 as a selective and reliable diagnostic, peripheral marker for AD, a very invaliding neurodegenerative condition that afflicts millions of people in the world, and for which none of the imaging or neurochemical diagnostic markers proposed to date can be considered to be sufficiently validated [50].

## Methods

### Purification of human neutrophils

Fresh human blood was from the Institute of Geriatric Rehabilitation “Santa Margherita” (Pavia, Italy) and from the Immunohematology and Transfusion Medicine Department of the “San Matteo” Hospital (Pavia, Italy) as approved by the respective internal Institution Review Boards. Neutrophils were obtained by Dextran-70 sedimentation of blood followed by Ficoll-Hypaque gradient centrifugation, essentially as described [51] with minor modifications [52], washed in PBS supplemented with 2 mM EDTA and 0.5% (w/v) BSA, and centrifuged at 500g for 10 min at 4 °C.

The residual erythrocytes in the neutrophils-rich fraction were eliminated by differential hypotonic lysis with ice cold 0.2% (w/v) NaCl for 30 s under agitation. The isotonicity was restored by adding an equal volume of 1.6% (w/v) NaCl. The cells were sedimented under the same conditions, and then washed three times in supplemented PBS. The number of neutrophils was determined by microscope-count, and the samples were then stored frozen at −80 °C.

### Determination of methionine sulfoxide reductase activity

Purified neutrophils were resuspended (10^8^ cells/ml) in PBS containing 1% (v/v) Triton X-100 and 0.1% (v/v) diisopropylfluorophosphate as a serine-protease inhibitor, and left on ice for 30 min. The lysate was then centrifuged at 18000 g for 10 min at 4°C. The supernatant was collected and analyzed directly. Protein concentration was assayed with the bicinchoninic acid method using BSA as a standard.

MsrB enzymatic activity was assayed using the *R* diastereoisomer of methionine sulfoxide conjugated with the dabsyl group as the substrate, and 1,4-dithioerythritol as the electron donor. Methionine-*R*-sulfoxide was separated from its optical antipode as described [53, 54] and functionalized with the dabsyl group as described [55]. The reaction mixture contained 5 mM sodium phosphate (pH 7.4), 154.5 mM NaCl, 4.5 mM KCl, 20 mM 1,4-dithioerythritol, 500 μM dabsyl-methionine-R-sulfoxide and 50 μg of total protein from the cell extract. The reaction was carried out at 37 °C for 30 min, and the reaction product, dabsyl-methionine, was analyzed by reverse-phase HPLC, monitoring the chromophoric dabsyl moiety at the wavelength of 436 nm [55].

## Acknowledgments

This work was supported by the University of Pavia, FAR funds to GM.

